# A Novel Xenonucleic Acid Mediated Molecular Clamping Technology for Early Colorectal Cancer Screening

**DOI:** 10.1101/2020.05.15.098954

**Authors:** Qing Sun, Larry Pastor, Jinwei Du, Michael J. Powell, Aiguo Zhang, Walter Bodmer, Jianzhong Wu, Shu Zheng, Michael Y. Sha

## Abstract

**Background:** Colorectal cancer (CRC) is one of the leading causes of cancer-related death. Early detection is critical to reduce CRC morbidity and mortality. In order to meet this need, we developed a molecular clamping assay called the ColoScape ™ for early colorectal cancer diagnostics.

**Methods:** Nineteen mutations in four genes APC, KRAS, BRAF and CTNNB1 associated with early events in CRC pathogenesis are targeted in the ColoScape™ assay. Xenonucleic Acid (XNA) mediated qPCR clamping technology was applied to minimize the wild-type background amplification in order to improve assay sensitivity of CRC mutation detection. The assay analytical performance was verified and validated, cfDNA and FFPE CRC patient samples were evaluated, and a ROC cure was applied to evaluate its performance.

**Results:** The data showed that the assay analytical sensitivity is 0.5% Variant Allele Frequency, corresponding to ~7-8 copies of mutant DNA with 5ng total DNA input per test. This assay is highly reproducible with intra-assay CV <3% and inter-assay <5%. We have investigated 380 clinical samples including plasma cfDNA and FFPE samples from patients with precancerous and different stages of CRC. The preliminary assay clinical specificity and sensitivity for CRC cfDNA were 100% (95% CI, 80.3-97.5%) and 92.2% (95% CI, 94.7-100%) respectively with AUC being about 0.96; and 96% (95% CI, 77.6-99.7%) specificity and 92% (95% CI, 86.1-95.6%) sensitivity with AUC 0.94 for CRC FFPE; and 95% specificity (95% CI, 82.5%-99.1%) and 62.5% sensitivity (95% CI, 35.8%-83.7%) with AUC 0.79 for precancerous lesions cfDNA.

**Conclusions:** XNA mediated molecular clamping assay is a rapid, precise, and sensitive assay for the detection of precancerous lesions cfDNA and CRC cfDNA or FFPE samples.

## BACKGROUND

Colorectal cancer (CRC) is one of the leading causes of cancer-related death and the third most common cancer with an estimated 1.8 million new cases worldwide in 2018 ^1^. Colorectal cancer usually starts as a noncancerous growth called an adenoma or polyp which takes many years until it may eventually develop into cancer ^2,3^. If detected at an early stage, CRC is treatable with a five-year survival rate of 90% while the survival rate drops to 12-14% if diagnosed at the advanced stage IV ^4^. Early detection is critical to reduce CRC morbidity and mortality.

There are several recommended CRC screening methods including colonoscopy and guiac-based fecal occult blood tests/fecal immunochemical tests (gFOBT/FIT) and FIT-DNA test ^2,5–7^. Colonoscopy is still the standard method for CRC screening but compliance with colonoscopy guidelines is low, possibly due to the invasive nature and the lengthy bowel preparation for the procedure, cost and potential complications during the procedure (6). gFOBT/FIT tests have been widely used in CRC screening; however, the test sensitivity and specificity are low ^8^. Sensitive and non-invasive methods for CRC screening are needed and advances in our understanding of molecular pathogenesis of CRC and molecular detection technologies now make this possible. Currently, non-invasive approaches include detection of genetic and epigenetic biomarkers associated with CRC in stool and plasma ^9–11^. Complex signaling pathways are involved in colorectal cancer pathogenesis, including WNT and RAS /RAF/MAPK pathways, microsatellite instability (MSI, DNA mismatch repair) and some gene specific CpG island methylation ^12–14^. Genetic and epigenetic changes in the pathways have been studied extensively in relation to their roles in the initiation and development of CRC ^15–28^.

KRAS mutations occur in 36-40% of CRC patients with majority of mutations at codons 12, 13 and 61 ^18,22,23^. The adenomatous polyposis coli (APC) gene is the key gene involved in the β-Catenin/Wnt signaling pathway and mutations in APC occur early and play an important role in colorectal tumorigenesis. The frequency of APC mutations ranges from 50% to 80% in CRC patients ^13,15–17^. BRAF is an oncogene which encodes a serine/threonine kinase that acts down-stream of KRAS in the MAPK pathway ^26–28^. BRAF mutations are present in about 10% of CRC with 90% of all BRAF mutations in CRC being BRAF V600E ^28^. Molecular characterization of CRC and data on mutation incidence in CRC provided the basis for biomarker selection for our CRC mutation testing. A panel of target genes APC, KRAS, BRAF, CTNNB1 was selected according to their mutation frequency in early stage colorectal cancer ^29–31^. One of the major challenges in cancer mutation detection is due to the fact that clinical samples from cancer patients frequently contain trace amounts of mutant allele in a large excess of wild-type DNA, which hampers sensitivity of mutation detection. Researchers have used different strategies to block or suppress wild-type effect on mutation detection, e.g. Cast-PCR, Cold PCR, ARMS ^32–34^ and blocking oligonucleotides employed in PCRs (e.g. 3’ spacer, 3’ phosphate, 3’ ddC, etc.) as well as nucleotides with unnatural backbones such as peptide nucleic acid (PNA) and locked nucleic acid (LNA) ^35,36^. However, all the mutation detection assays employing the strategies and methods currently available still have limited sensitivity for detection of low-abundance variants, especially at early stage cancer, when mutations are present in less than 1% VAF or even much lower ratios of mutant to wild-type sequence. In this study, we applied molecular Clamping technology ^37,38^ for minimally invasive and sensitive detection of CRC mutations from liquid biopsy or tumor tissue. This technology is employed to suppress amplification of wildtype alleles and so improve the sensitivity of mutation detection, especially for early adenomas and early stage CRC. Xenonucleic acid (XNA) is a synthetic DNA analog in which the phosphodiester backbone has been replaced by a novel synthetic modified backbone chemistry (Figure 1a). XNA’s are highly effective at hybridizing to targeted normal DNA sequences and can be employed as molecular clamps in quantitative real-time polymerase chain reactions (PCR) or as highly specific molecular probes for detection of nucleic acid target sequences 39 (Roberta D’Agata 2017). Binding of XNA to its target sequence blocks strand elongation by DNA polymerase in PCR assays. When there is a mutation in the target site, and therefore a mismatch, the XNA-DNA duplex is unstable, allowing strand elongation by DNA polymerase. XNA oligomers also are not recognized by DNA polymerases and cannot be utilized as primers in subsequent real-time PCR reactions (Figure 1b).

**Figure 1.**
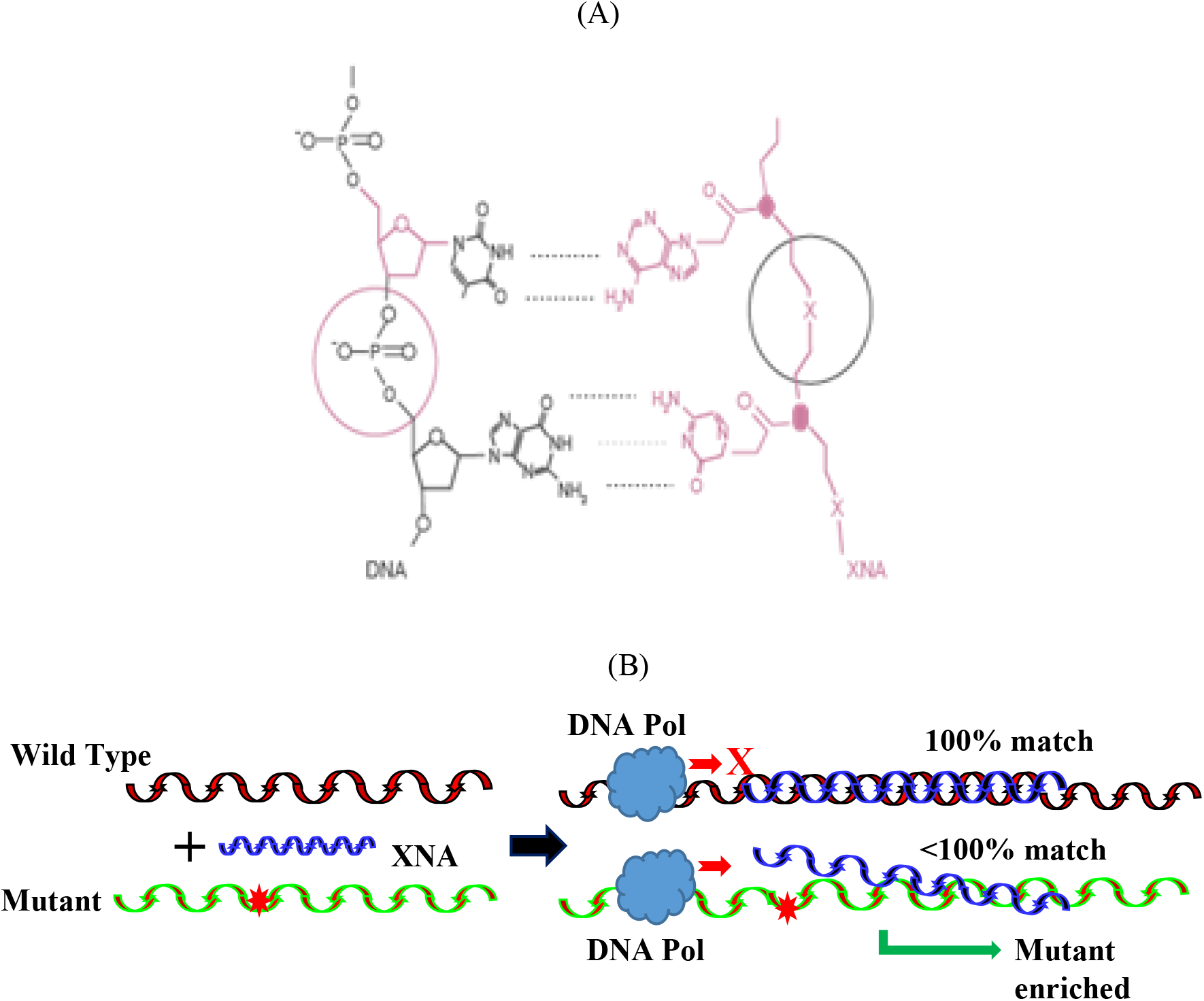
XNA structure and its function in the assay. (A). XNA structure and hybridization with DNA. (B). Principle of the Coloscape™ mutation detection in targeted genes. XNAs hybridize tightly to complementary DNA target sequences only if the sequence is a complete match. When there is a mutation in the target site, and therefore a mismatch, the XNA: DNA duplex is unstable, allowing strand elongation by DNA polymerase. Addition of an XNA, whose sequence is a complete match to the wild-type DNA, to a PCR reaction, blocks amplification of wild-type DNA allowing selective amplification of mutant DNA. This enrichment of the mutation amplicons enables mutation detection by qPCR.

Herein we report the development and validation of a novel XNA-based multiplex real-time PCR assay for simultaneous and qualitative detection of somatic mutations in the genes frequently mutated in CRC patients. This multigene biomarker assay, called ColoScape™, includes target gene mutation detection in APC, KRAS, BRAF, and CTNNB1 (Table 1).

**Table 1.** List of Coloscape™ Targeted Gene Mutations

| Genes | Exon | Amino Acid Change | Nucleotide change | Cosmic No. |
| --- | --- | --- | --- | --- |
| KRAS | 2 | p.G12>A | c.35G>C | COSM522 |
|  |  | p.G12>R | c.34G>C | COSM518 |
|  |  | p.G12>D | c.35G>A | COSM521 |
|  |  | p.G12>C | c.34G>T | COSM516 |
|  |  | p.G12>S | c.34G>A | COSM517 |
|  |  | p.G12>V | c.35G>T | COSM520 |
|  |  | p.G13>D | c.38G>A | COSM532 |
|  |  | p.G13>C | c.37G>T | COSM527 |
|  |  | p.G13>R | c.37G>C | COSM529 |
| APC | 15 | p.E1309fs* | c.3921_3925delAAAAG | COSM18764 |
|  |  | p.Q1367* | c.4099C>T | COSM13121 |
|  |  | p.R1450* | c.4348C>T | COSM13127 |
|  |  | p.R876* | c.2626C>T | COSM18852 |
| CTNNB1 | 3 | p.T41A | c.121A>G | COSM5664 |
|  |  | p.T41I | c. 122C>T | COSM5676 |
|  |  | p.S45P | c.133T>C | COSM5663 |
|  |  | P.S45F | c.134C>T | COSM5667 |
|  |  | P.S45del | c.133-135delTCT | COSM6128 |
| BRAF | 15 | p.V600E | c.1799T>A | COSM476 |

## MATERIALS AND METHODS

### Reference materials and clinical samples

The following genomic DNA reference materials carrying specific mutations were obtained from ATCC and Horizon Discovery Group plc respectively: CTNNB1 S45 (HCT116), APC E1309 (LS1034), APC Q1367 (C2BBe1), APC R1450 (SW837), KRAS G12 (Horizon Cat#: HD272), KRAS G13 (Horizon Cat#: HD290), BRAF V600 (Horizon Cat#: HD238). For target mutations which no commercial reference materials were available, APC, CTNNB1 synthetic DNA templates from Integrated DNA Technologies Inc were used. Reference cfDNA standards APC R1450, CTNNB1 T41, KRAS G12 and BRAF V600E were purchased from SeraCare Inc.

For reference cfDNA standards that are not available commercially, genomic DNAs carrying APC E1309/Q1367, CTNNB1 S45 and KRAS G13 mutations were sheared by sonication with M220 Focused-ultrasonicator (Covaris Inc). The sonicated DNAs were analyzed on BioAnalyzer (Agilent) to give an average DNA fragment length of about 150 bp which mimics the size of cfDNA fragments justifying their use as cfDNA references.

Most of FFPE and plasma (cfDNA) clinical samples with CRC used for this study were collected from Chinese patients (Second Affiliated Hospital of Zhejiang University, Hangzhou and Jiangsu Cancer Hospital, Nanjing, China). The ethics approval was awarded by Ethics Committee of the Second Affiliated Hospital of Zhejiang University, Hangzhou, China; and Ethics Committee of Jiangsu Cancer Hospital & Jiangsu Institute of Cancer Research, Nanjing, China). All subjects provided written informed consent. 10 mL blood were drawn from each patient and stored in cfDNA BCT Streck tubes.

### DNA Extraction

DNA from FFPE samples was extracted with the QIAamp DSP DNA FFPE Tissue Kit (Catalog, Qiagen, REF 60604. QIAGEN GmbH, Hilden, Germany) following manufacturer’s instructions. For cfDNA isolation, the collected blood was first spun at 1600xg on a table centrifuge (Sorvall ST16R, Thermo Fisher Scientific) for 10 minutes at room temperature. The supernatant above the interface phase was carefully taken and spun at 16,000xg for 10 minutes at room temperature. The final plasma supernatant was stored at −20 °C until use. cfDNAs were isolated from plasma by using QIAamp ® MinElute ccfDNA Midi Kit (QIAGEN, Cat# 55284) following the manufacturer’s instructions. Isolated cfDNA was quantified by using Qubit dsDNA HS Assay Kit (Thermo Fisher Scientific, Cat # Q32851) and also assessed by using the beta-actin qPCR assay (as internal control) to check the quantity and quality.

### qPCR Primer and Probe Design

The high sensitivity of this multigene biomarker assay is achieved due to XNA clamp probe technology. XNA oligomers are designed that bind to the selected wild-type sequences at the respective genetic loci in the target genes. For each of the selected mutation sites, primers and TaqMan hydrolysis probes were designed by Primer3 software version 0.4.0. For target gene codons with multiple mutations, e.g. KRAS G12 six mutations, a locus specific probe was designed so that all 6 of KRAS G12 mutations can be detected in one assay using one pair of primers with the same XNA designed for the relevant position in KRAS G12. For target gene codons with single mutations, e.g. APC E1309, APC Q1367 and APC R1450 and APC R876, mutant specific probes or allele specific primers were designed. The human beta actin gene (ACTB) was selected as an internal control for the assay (Supplementary Table 1). The designed primers and probes were analyzed in silico to verify the specificity of the oligos (by GenBank Blast against a whole genome reference DNA), no primer dimers (Auto-Dimer), no amplicon secondary structure (M-fold) before synthesis. All primers were synthesized by IDT (Integrated DNA Technology) and probes were ordered from BioSearch Inc.

### XNA Synthesis

XNA was synthesized in house.

### ColoScape ™ Assay

The assay consists of 10 μl of reaction volume including 5 ul of 2X buffer (Bioline, Bio-11060), 2 ul of Primer/probe mix in 1xTE with final concentration of 100 nM-600 nM of primers and 50 nM – 500 nM of probes, 1 ul of XNA final concentration from 0.125 μM to 1 μM and 2 μl of template (nuclease free water for non-template control or 5-10 ng DNA). Non-template controls (NTC), Clamping controls (CC, human wildtype gDNA) and positive controls (PC, include each mutant DNA) were included in each run. The thermocycling profile is as follows: 95 °C for 2 minutes followed by 50 cycles of 95 °C for 20 seconds, 74 °C for 40 seconds, 62 °C for 30 seconds and 72 °C for 30 seconds. The assay consists of three multiplex qPCR reactions with XNAs to simultaneously detect all the indicated mutations (Supplementary Table 2).

The mutational status of a sample was determined by calculating the Cq value between amplification reactions for a mutant allele assay and an internal control assay. Cq difference (ΔCq) = Mutation Assay Cq - Internal Control Assay Cq. The cut-off values were experimentally determined as its ΔCq value by testing at least 20 wildtype gDNA and/or cfDNA repeatedly during the verification of assay performance. Cut-off ΔCq is calculated as ΔCq cut-off =ΔCq Cq average – 1.96*SD (at 99% CI). If the sample ΔCq is ≤ cut-off value, the mutation is detected as positive. If the sample ΔCq is > cut-off value, the mutation is not detected.

### Performance parameters of the assay

Performance parameters of the assay were established on DNA samples extracted from FFPE and plasma of CRC patients as well as reference materials. Assay performance characteristics were verified with respect to precision, limit of detection, specificity and cross-reactivity as well as clinical sample validation and comparison with Sanger Sequence or NGS.

### Statistical analysis

We calculated the sensitivity, specificity, and Area Under Curve (AUC) of each group by sklearn.metrics ^40^. ROC curves were then plotted by Matplotlib. Pyplot ^41^. ROC curve and the area under the curve (AUC) were used to describe the assay performance.

## RESULTS

### Assay Feasibility

We designed the primers and probes for APC, CTNNB1, BRAF and KRAS, plus XNAs that cover 19 mutations of these 4 genes. To demonstrate that XNA can effectively suppress wild-type allele amplification and thus enrich the mutations detection during qPCR, we compared qPCR with and without XNA. The data show that XNA-based qPCR has delta Ct ~9 for mutant to wildtype (Figure 2A) whereas qPCR without XNA has delta Ct ~0.3 for mutant to wildtype (Figure 2B). Sanger sequencing also confirms that amplicons from XNA-based qPCR have a pure mutation reading (GCC for CTNNB1 T41A, Figure 2C) while amplicons from qPCR without XNA have a mixed reading of wildtype and mutation (A/GCC for CTNNB1 T41A, Figure 2D). XNA-based qPCR for other target gene mutants also show the same pattern except BRAF V600E (Figure 2E). This demonstrates that XNA enables mutation detection easily by blocking wildtype sequence amplification. For BRAF V600E, an allele specific primer was designed to genotype BRAF V600E directly (Figure 2E_a). To verify that assay sensitivity was not compromised by multiplexing, we also compared singleplex and multiplex qPCR for each gene mutation in 1% VAF reference DNA sample. The data show that multiplex qPCR for each gene mutation has almost identical Cq value when compared to that of singleplex qPCR (Supplementary Table 3).

**Figure 2.**
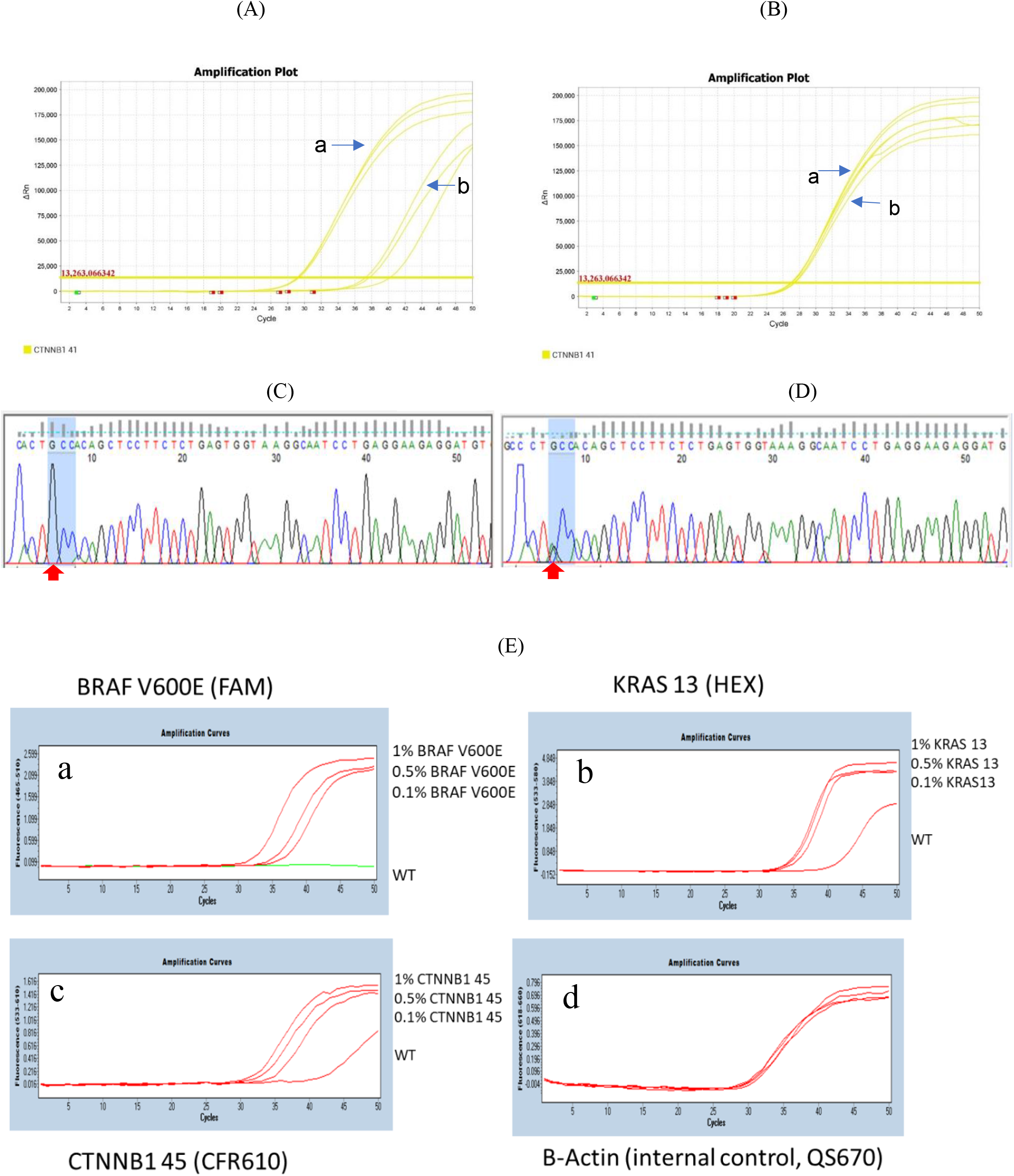
Comparison of XNA-based qPCR and without XNA qPCR. (A). CTNNB1 T41 amplification curves with XNA. a: 5% VAF of CTNNB1 T41 mutant, CT = 29.1 (ACTB, CT = 30.7, unshown here due to use Cy5 channel), b: CTNNB1 T41 wildtype, CT = 38.3 (ACTB, CT = 30). It indicates that XNA-based qPCR has delta Ct ~9 for mutant to wildtype. (B). CTNNB1 T41 amplification curves without XNA. a, 5% VAF of CTNNB1 T41 mutant, CT = 26.9 (ACTB, CT = 30.4). b, CTNNB1T41 wildtype, CT = 27.2 (ACTB, CT = 30). It indicates that qPCR without XNA has delta Ct ~0.3 for mutant to wildtype. (C). Sanger Sequencing for amplicon from CTNNB1 T41 assay with XNA, it confirms that there is only mutant GCC at CTNNB1 T41 (red arrow). (D). Sanger Sequencing for amplicon from CTNNB1 T41 assay without XNA. it shows there is mix of wildtype and mutant A/GCC of CTNNB1 T41(red arrow). (E). Amplification profile of ColoScape™ multiplex qPCR assay with various concentrations of reference gDNA. a, BRAFV600E with 1%, 0.5%, 0.1% and 0% VAF (Fam as probe labeling dye). b. KRAS G13 with 1%, 0.5%, 0.1% and 0% VAF (Hex as probe labeling dye); c. CTNNB1 S45 with 1%, 0.5%, 0.1% and 0% VAF (CFR610 as probe labeling dye); and d. Beta-Actin (internal control, QS670 as probe labeling dye).

### The Assay Analytical Sensitivity

The analytical sensitivity of the ColoScape™ were determined by studies involving APC, CTNNB1, KRAS and BRAF-defined genomic DNA reference samples. These known variant allele reference samples were diluted to 1% VAF (variant allele frequency), 0.5% VAF and 0.1% VAF separately. The reference samples at 5ng and 10 ng input were evaluated. For all tested purified reference cfDNA inputs from 5 - 10 ng/well, all target mutations were detected with 100% correct calls at 0.5% VAF (Table 2) so the overall limit of detection (LOD) is 0.5% VAF. Moreover, sensitivity testing for APC Q1367, APC R1450, CTNNB1 T41, KRAS G12 &G13 and BRAF V600 in plasma even showed 0.1% VAF detection at 5 ng DNA (Table 2). For FFPE gDNA samples, the LOD for this ColoScape™ assay is overall 0.5% VAF at 5 ng DNA input which is about ~7-8 copies of mutant DNA (1 ng gDNA about 330 genomic copies). This was confirmed in three qPCR instruments ABIQS5, ABI 7500 FAST Dx and Roche LC480II (Supplementary Table 4).

**Table 2.**
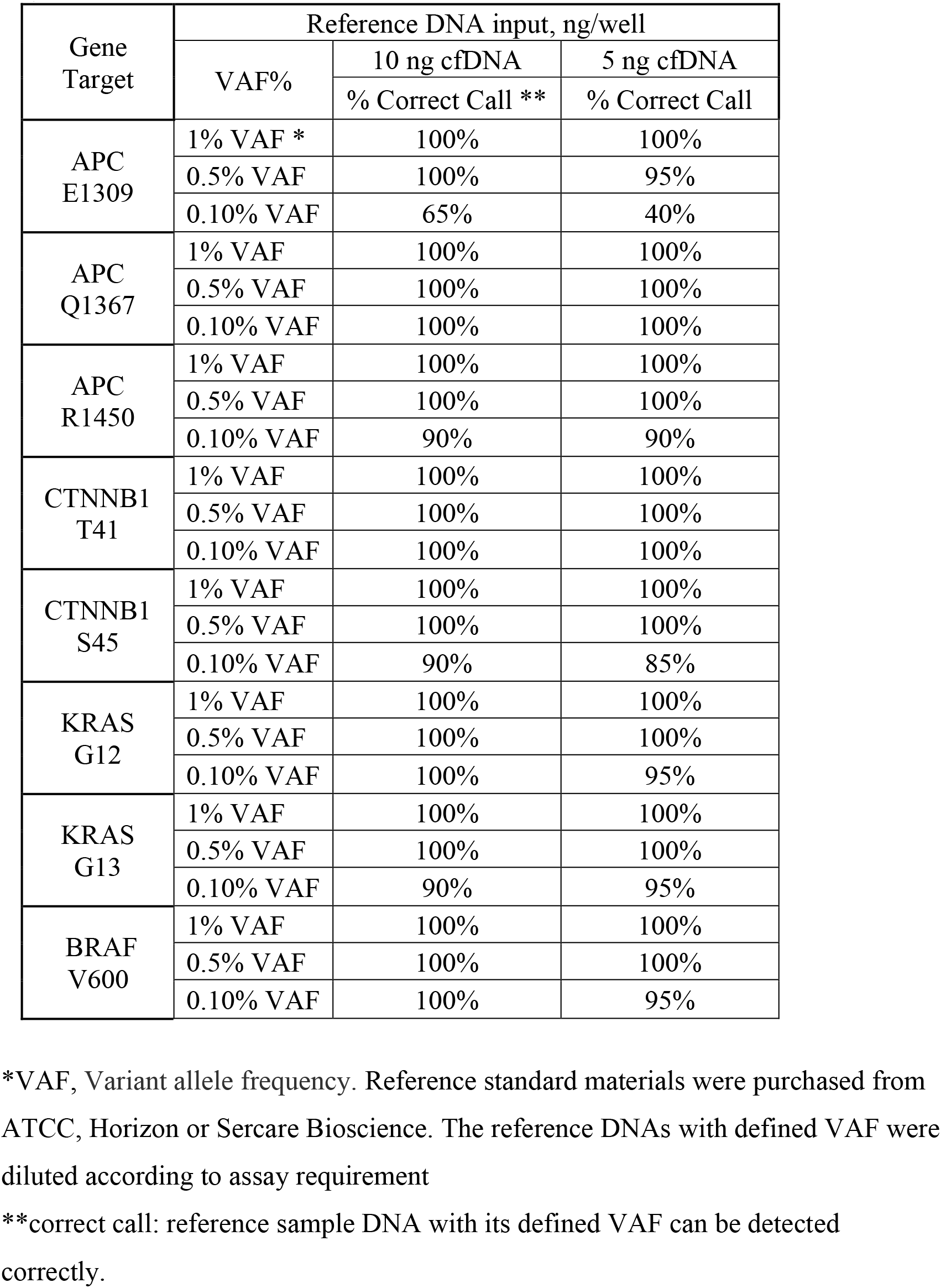
Summary of Assay Limit of Detection for cfDNA Sample

| Gene Target | Reference DNA input, ng/well |  |  |
| --- | --- | --- | --- |
|  | VAF% | 10 ng cfDNA | 5 ng cfDNA |
|  |  | % Correct Call ** | % Correct Call |
| APC E1309 | 1% VAF * | 100% | 100% |
|  | 0.5% VAF | 100% | 95% |
|  | 0.10% VAF | 65% | 40% |
| APC Q1367 | 1% VAF | 100% | 100% |
|  | 0.5% VAF | 100% | 100% |
|  | 0.10% VAF | 100% | 100% |
| APC R1450 | 1% VAF | 100% | 100% |
|  | 0.5% VAF | 100% | 100% |
|  | 0.10% VAF | 90% | 90% |
| CTNNB1 T41 | 1% VAF | 100% | 100% |
|  | 0.5% VAF | 100% | 100% |
|  | 0.10% VAF | 100% | 100% |
| CTNNB1 S45 | 1% VAF | 100% | 100% |
|  | 0.5% VAF | 100% | 100% |
|  | 0.10% VAF | 90% | 85% |
| KRAS G12 | 1% VAF | 100% | 100% |
|  | 0.5% VAF | 100% | 100% |
|  | 0.10% VAF | 100% | 95% |
| KRAS G13 | 1% VAF | 100% | 100% |
|  | 0.5% VAF | 100% | 100% |
|  | 0.10% VAF | 90% | 95% |
| BRAF V600 | 1% VAF | 100% | 100% |
|  | 0.5% VAF | 100% | 100% |
|  | 0.10% VAF | 100% | 95% |

\*VAF, Variant allele frequency. Reference standard materials were purchased from ATCC, Horizon or Sercare Bioscience. The reference DNAs with defined VAF were diluted according to assay requirement
\*\*correct call: reference sample DNA with its defined VAF can be detected correctly.

### The Assay Precision

The high precision of the assay was verified by testing inter- and intra-assay, lot-to-lot (3 different lots), operator and instrument variability. Instruments tested produced consistent results for the 1% mutant and wildtype gDNA controls. Ct values from three replicates were averaged and standard deviation and coefficient of variation (CV) were calculated. The precision studies indicate that this assay is highly reproducible with intra-assay CV <3%, inter-assay CV <5%, lot-to-lot CV <4% and operator variability CV <3%. Therefore, the assay analytical precision is high.

### The Assay Analytical Specificity

With known reference wildtype gDNA, the assay specificity is over 97%. There were no false positive calls for up to 320 ng of gDNA per reaction and up to 20ng FFPE DNA per reaction with results confirmed by NGS. Overall, the analytical specificity of the assay was over 95% on reference gDNA and DNA extracted from FFPE or plasma.

Another aspect of the assay specificity is manifested by evaluation of assay cross-reactivity. Each target assay was tested against all positive reference material to evaluate the cross-reactivity. All target mutations except KRAS G12 were detected as expected by the multiplex ColoScape™ (Table 3), indicating there is no cross-reactivity for these different target detections. KRAS G12 produced a signal in KRAS G13 positive samples. However, there is a 6 Ct difference between the true KRAS G13 signal and the crosstalk signal from KRAS G12. Furthermore, since the kit is to detect KRAS G12 and KRAS G13 mutations separately in different tubes, the crosstalk will not have any impact on the assay performance. Therefore, only intended target mutations can be detected by the ColoScape™ assay.

**Table 3.**
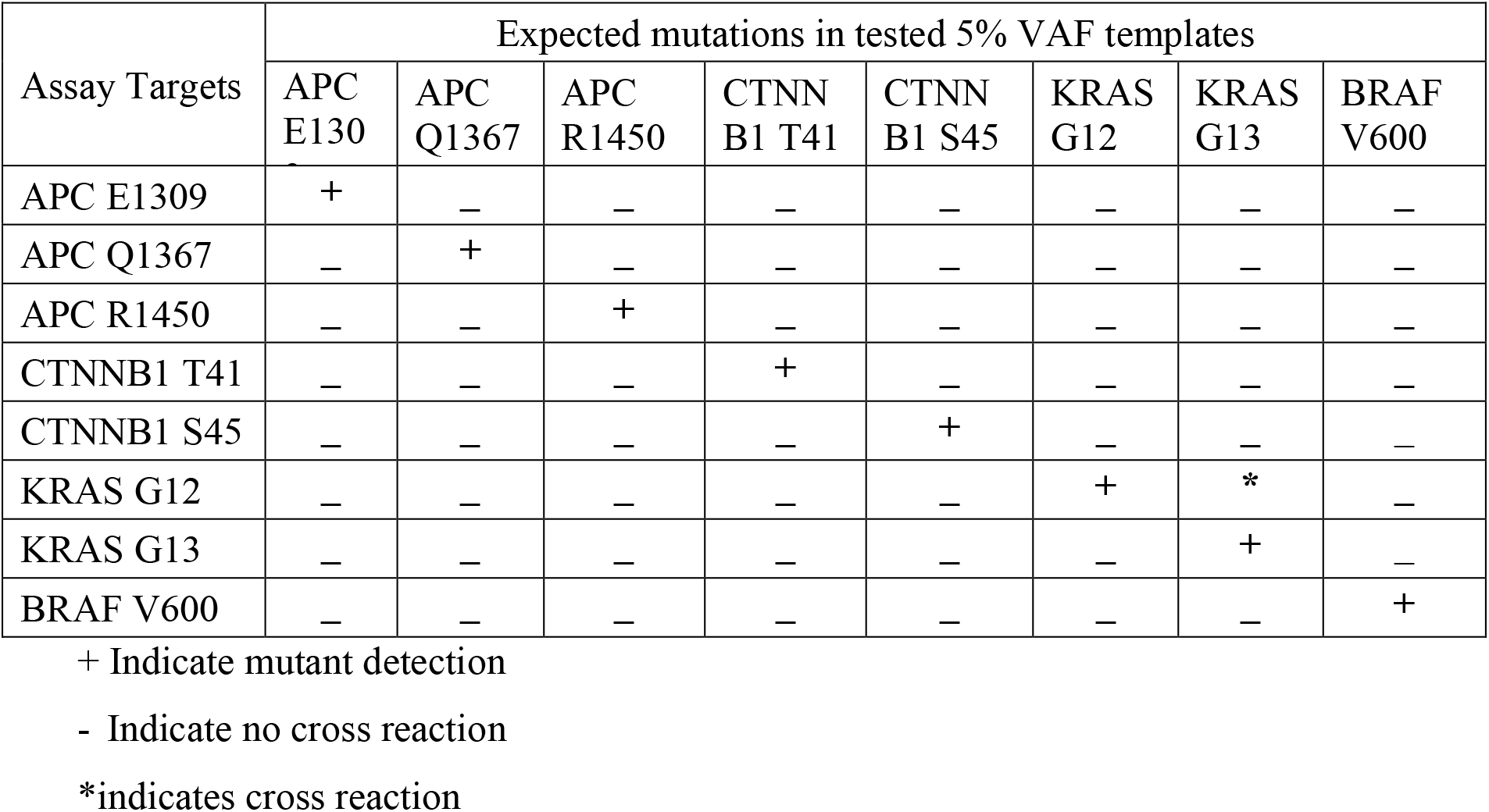
Assay Analytical Specificity – the Assay Cross Reactivity

| Assay Targets | Expected mutations in tested 5% VAF templates |  |  |  |  |  |  |  |
| --- | --- | --- | --- | --- | --- | --- | --- | --- |
|  | APC E130 | APC Q1367 | APC R1450 | CTNN B1 T41 | CTNN B1 S45 | KRAS G12 | KRAS G13 | BRAF V600 |
| APC E1309 | + | – | – | – | – | – | – | – |
| APC Q1367 | – | + | – | – | – | – | – | – |
| APC R1450 | – | – | + | – | – | – | – | – |
| CTNNB1 T41 | – | – | – | + | – | – | – | – |
| CTNNB1 S45 | – | – | – | – | + | – | – | – |
| KRAS G12 | – | – | – | – | – | + | * | – |
| KRAS G13 | – | – | – | – | – | – | + | – |
| BRAF V600 | – | – | – | – | – | – | – | + |

+ Indicate mutant detection
- Indicate no cross reaction
\*indicates cross reaction

### Clinical Performance

Clinical sensitivity and specificity were evaluated on FFPE and plasma of patients with different stages of CRC from normal to advanced adenomas (AA), to CRC stages I through IV. 380 patient samples including 185 FFPE and 195 cfDNA were tested during different experiment periods. In each case, a sample was considered positive if at least one of the target mutations tested positive. The test result was considered correct if the CRC samples and health samples were confirmed by Sanger sequencing or NGS.

To comparing ColoScape ™ and NGS, eighteen cfDNA samples of CRC patients were tested with ColoScape ™ and NGS. The data show that ColoScape ™ and NGS has a concordance rate of 89 % (Table 4). There were 2 samples (DC04 and DC07) identified as mutants by ColoScape and confirmed by Sanger sequencing, but not detectable by NGS.

**Table 4.** Comparison of ColoScape ™ and NGS for CRC cfDNA Samples

| Patient ID | ColoScape Results | NGS Results |
| --- | --- | --- |
| DC01 | Negative | Negative |
| DC02 | KRAS G12 Positive | KRAS G12C |
| DC03 | KRAS G12 Positive | KRAS G12D |
| DC04 | KRAS G12 Positive | Negative |
| DC05 | KRAS G12 Positive | KRAS G12C |
| DC06 | APC E1309/1367 Positive, KRAS 12 Positive | APC E1309FS* |
| DC07 | KRAS G12 Positive | Negative |
| DC08 | KRAS G13 Positive | KRAS G13 |
| DC09 | KRAS G12 Positive | KRAS G12C |
| DC10 | KRAS G12 Positive | KRAS G12 |
| DC11 | Negative | Negative |
| DC12 | Negative | Negative |
| DC13 | KRAS G12 Positive | KRAS G12D |
| DC14 | APC R1450/876 Positive, KRAS G12 Positive | KRAS G12D |
| DC15 | KRAS G13 Positive | KRAS G13D |
| DC16 | KRAS G12 Positive | KRAS G12D |
| DC17 | KRAS G12 Positive | KRAS G12C |
| DC18 | KRAS G13 Positive | KRAS G13D |

We also compared ColoScape ™ and Sanger Sequencing (amplicon of qPCR) of 97 FFPE samples, which showed that the ColoScape ™ has 98% concordant rate with the Sanger sequencing (Supplementary Table 5).

To investigate whether plasma cfDNA and FFPE differ for CRC mutations detection from same patient, we tested 11 pairs of matching tumor FFPE/adjacent normal tissues and cfDNA samples covering CRC stages I-IV. (Table 5). Interestingly, there were 100% cfDNA and 91% FFPE shown detectable mutations. Its concordance for FFPE and cfDNA about 90% (10/11 samples). There was one patient identified with different mutant for their cfDNA and FFPE paired samples (DCS9).

**Table 5.**
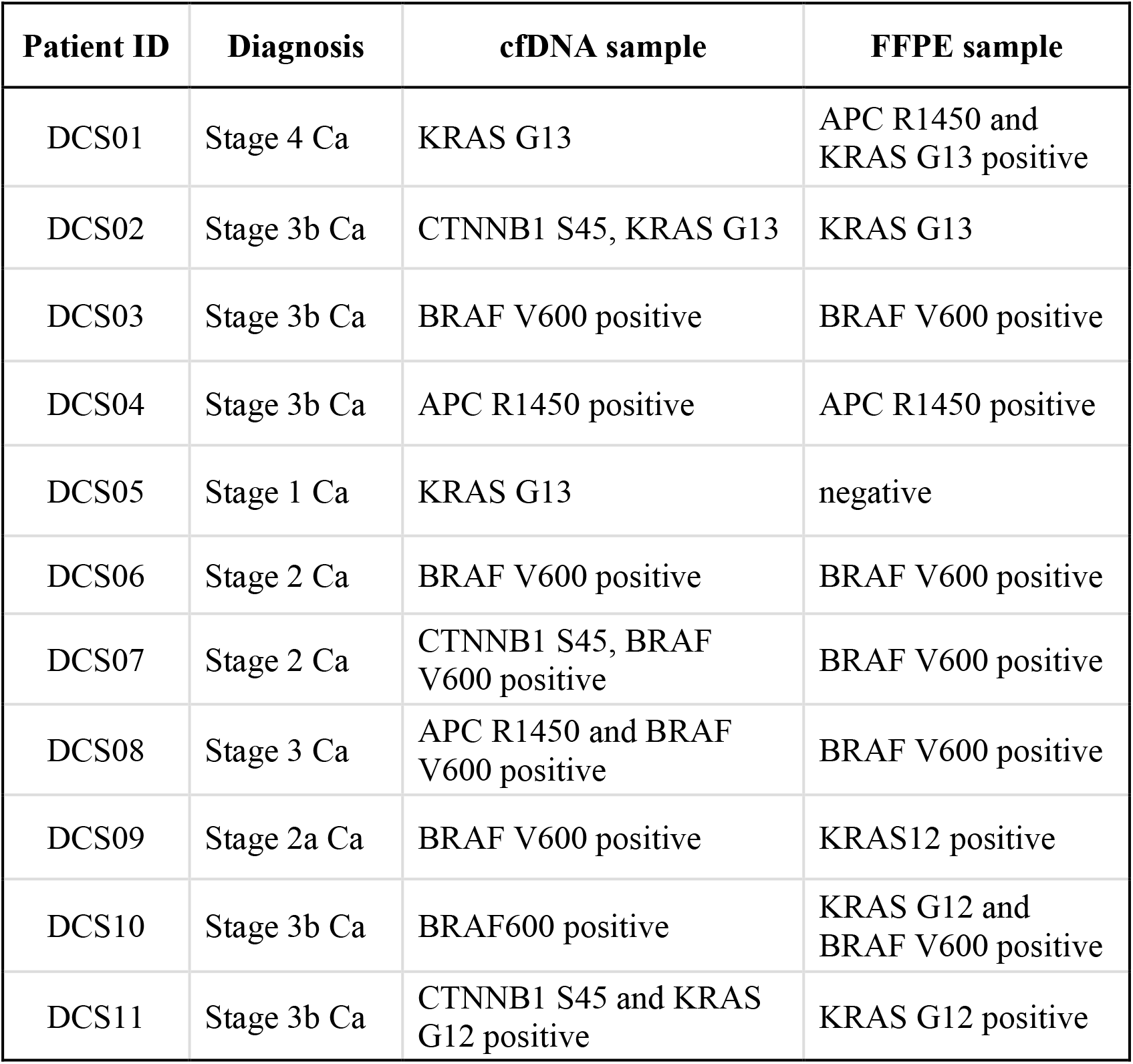
Comparison of Pairs of cfDNA and FFPE CRC Patient Samples

For precancerous screening, we tested 10 advanced adenomas samples (AA, FFPE) and detected 6 true positives and 3 false negatives and confirmed its sensitivity at 66.6 % (Table 6). For precancerous screening with cfDNA, 58 plasma samples from FIT + patients were tested by this assay and colonoscopy. There were 2 false positives with 40 true negative samples, yielding a specificity of about 95.2%. There were 6 false negatives with 10 true positive samples which were confirmed by Sanger sequencing, corresponding to sensitivity of about 62.5% for this cfDNA precancerous screening. This preliminary data indicates that this assay has a 62.5% - 66% sensitivity for precancerous screening (AA FFPE 66.6% [95% CI, 30.9 %-90.9%]; and cfDNA 62.5% [95% CI, 35.8%-83.7%]) and specificity for precancerous cfDNA was 95.2% (95% CI, 82.5%-99.1%), its AUC about 0.79 (Table 6, Figure 3a). Excluding precancerous screening samples, for CRC FFPE, sensitivities 92% (95% CI, 86.1%-95.6%) and specificities were 96% (95% CI, 77.6%-99.7%), its AUC about 0.94 (Table 6, Figure 3b); while for CRC cfDNA, sensitivities were 92.2% (95% CI, 80.3%-97.5%) and specificities were 100% (95% CI, 94.7%-100%), its AUC 0.96 (Table 6 and Figure 3c). This assay accuracy is about 92.5% for CRC FFPE and 97% for CRC cfDNA separately (Table 6).

**Table 6.** Clinical Sensitivity and Specificity for FFPE and cfDNA Samples

| **Types of Clinical Samples** | **Patient number** | **Specificity\*** | **Sensitivity** | **Accuracy** | **PPV** | **NPV** |
| --- | --- | --- | --- | --- | --- | --- |
| CRC FFPE | 175 | 96.0% | 92.0% | 92.5% | 99.20% | 66.60% |
| CRC cfDNA | 137 | 100.0% | 92.2% | 97.0% | 100% | 95.50% |
| Advanced Precancerous lesions (AA *, FFPE) | 10 | NA | 66.60% | 66.6% | 100% | NA |
| Precancerous lesions (cfDNA) | 58 | 95.20% | 62.50% | 86.2% | 83.30% | 86.90% |

\* AA, Advanced Adenomas

**Figure 3.**
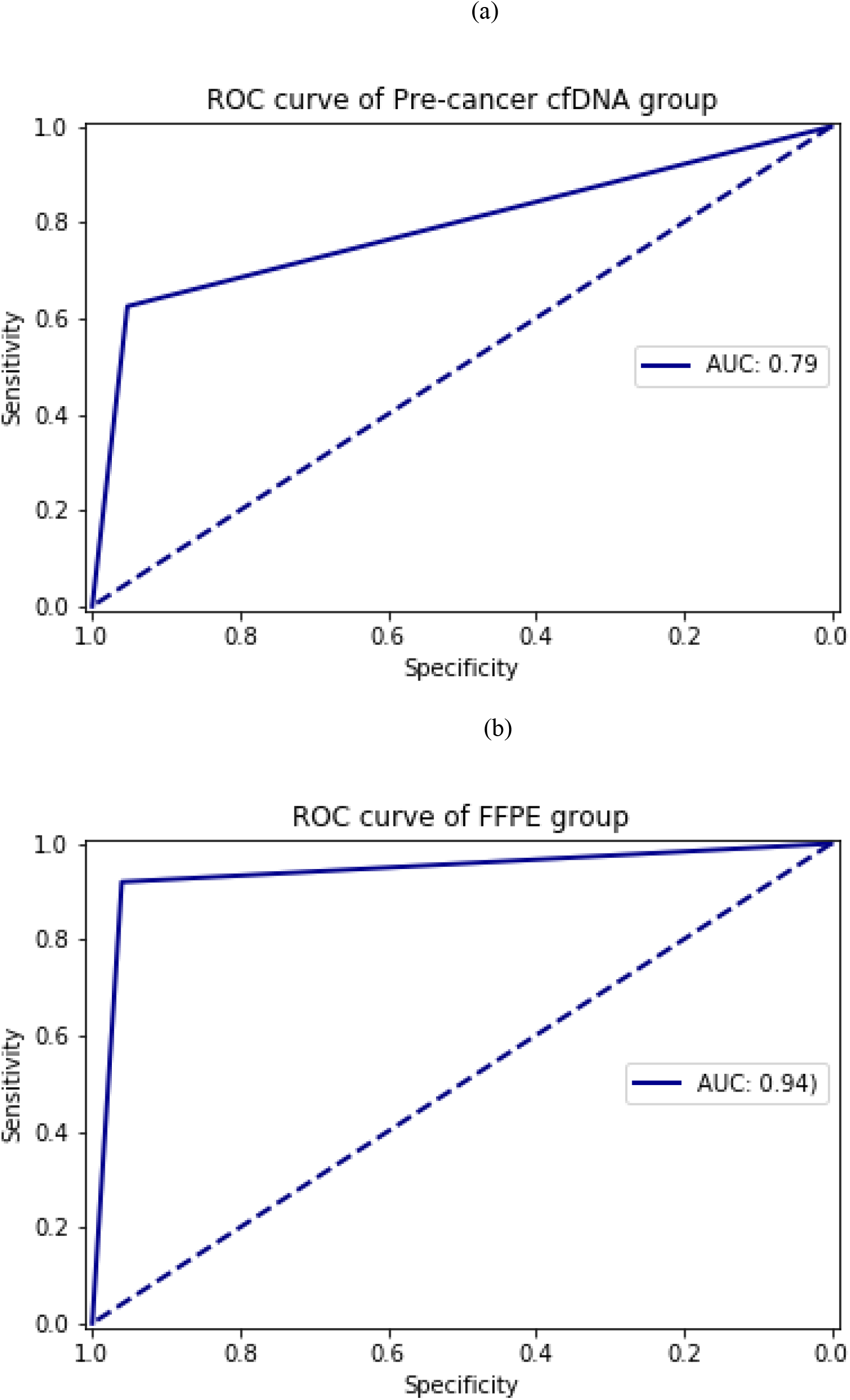
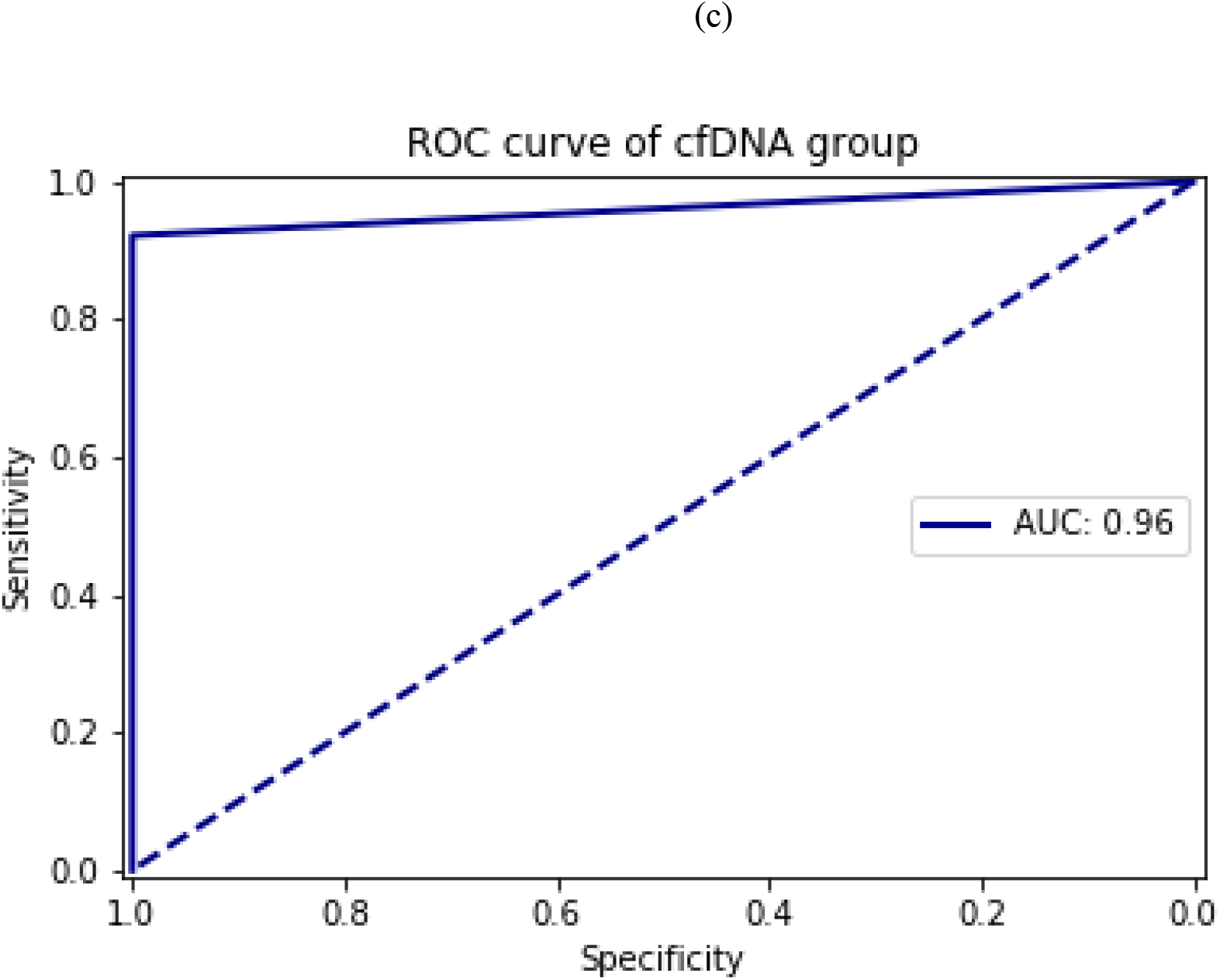
Receiver operating characteristic (ROC) curve and area under the curve (AUC) to evaluation the performance of the assay. a precancerous lesions cfDNA samples analysis. b, CRC FFPE samples analysis; c, CRC cfDNA samples analysis.

## DISCUSSION

In this study, ColoScape ™ was shown to be a robust CRC mutation detection assay with analytical sensitivity (LOD) about 0.5% VAF for cfDNA and FFPE. For some targets, their LOD was up to for 0.1%VAF with preliminary clinical sensitivity about 92% and specificity about 96-100%, however, this needs further clinical trial for confirmation. The ColoScape ™ assay is based on xenonucleic acid (XNA) mediated molecular clamping technology. XNA has a novel unique chemical backbone which is distinguished from the ‘classical’ PNA by replacement of the backbone methylene functionalities by a heteroatom which endows XNA molecular clamps with high binding affinity for both DNA and RNA templates and higher melting temperature differentials in SNV and indels against natural DNA. The effectiveness of XNA in suppressing wild type amplification has also been demonstrated in mutation detection using a ‘classical’ PNA PCR Clamp detection assay or by Sanger sequencing or NGS ^35,36^. XNA is thus confirmed to be a novel oligo blocker that will be applicable to a variety of cancer mutation detection assays to improve assay sensitivity. Araki et al reported that PNA-Clamp SmartAmp2 which its principle is similar as ColoScape assay, can detect as little as 1% of the mutant allele of KRAS G12 in the DNA samples ^42^. However, our assay can detect KRAS G12 even as lower as 0.1% VAF from cfDNA sample (Table 2). Another advantage of XNA is that mutations can be detected that are in the genomic region covered by XNA. As shown in this study, only one pair of primers and probe are needed in the assay for detection of all 6 genotypes of KRAS G12 mutations, which makes multiplexing of more target mutation detection easier due to the reduced need for PCR primers and probes and so reduced competition among reaction components. The LODs of different variants range from 0.2% to 2.5% VAF (Supplementary Table 6), possibly due to variations in the nature of primer-template mismatches. This is consistent with observations on the effects of primer-template mismatches on the detection and quantification of nucleic acids ^43^.

The preliminary clinical performance of the ColoScape ™ assay showed good sensitivity, including the testing of precancerous screening samples. Mutations from early cancer patients were detected by ColoScape ™ and confirmed by Sanger sequencing. ColoScape and Sanger Sequencing were 98% concordant. ColoScape has about 89% concordance with NGS (Table 4). There were two samples identified as mutants by ColoScape and confirmed by Sanger sequencing, but not detectable by NGS, possibly due to higher assay sensitivity (LOD about 0.1% VAF) of ColoScape than that of NGS (LOD about 1% VAF). ColoScape has been used in testing of precancerous plasma from FIT positive patients in a small study and showed a positive predictive value (PPV) of 83.3% and assay sensitivity 62.5% while FIT testing has a PPV of 25%. Since FIT has such a high false negative rate, potentially, ColoScape as a triage test in combination with a FIT test will improve the effectiveness of CRC patient management and treatment. There are several CRC-related mutation detection and methylation based screening assays, e.g. Cologuard (Exact Sciences Corporation), Epi ProColon® (Epigenomics AG), However, these FDA approved assays are based on only a single gene methylation detection (Septin 9-Epi ProColon) or 2 methylation biomarkers combined with only one target gene mutation assay (Cologuard) which potentially limits assay sensitivity. In this study we have preliminary shown that ColoScape ™ can detect 66.6% of mutations from advanced adenoma (AA) samples although its only 10 samples which needs further confirmed, while the reported Cologuard AA detection rate is 42-46% ^44, 45^. This suggests that ColoScape ™ potentially has comparable sensitivity for early colon cancer detection from patient’s blood samples. Furthermore, we have demonstrated a sensitivity for cfDNA sample for precancerous lesions about 62.5% with specificity 95.2% which it has great potential for early precancerous lesions screening.

We have shown that the ColoScape ™ assay has a robust analytical performance and preliminary clinical accuracy for FFPE and plasma samples (Table 2 &5, Supplementary Table 5). This rapid, precise and sensitive molecular assay for mutation detection in CRC has the following key benefits: (a) The assay is unique. It covers a selection of multiple clinically relevant mutations in 4 genes and the proprietary XNA QClamp® TaqMan-based PCR technology has an increased mutation detection sensitivity. (b). The assay is easy to use. Multiplex qPCR assay enables easy assay setting up by end users. (c). The assay is efficient. Only 15-30 ng DNA is needed for assay input. (d). The assay is specific & sensitive. No cross-reactivity with wild-type even with up to 320 ng purified gDNA. With 10ng cfDNA or FFPE DNA per reaction, mutations can be detected at 0.5% VAF and even up to 0.1% VAF. (e) Preliminary assay clinical specificity is about 95-100% depending on sample type (f). The assay reaction is rapid. Total run time is less than 3 hours. (g). The assay is versatile: assay validated on widely used real-time qPCR machines. This open instrument system is more convenient for different clinical laboratories end users.

In summary, we have developed a rapid and sensitive assay to enable molecular characterization and detection of precancerous and different stages of CRC in a variety of samples. However, this XNA-based qPCR with only five channels available in qPCR instrument has its limitations for which the assay can be improved by targeting more genes and more mutation hotspots. Recent molecular characterization and NGS analysis of cancer patients has provided unprecedent insight into cancer molecular mechanisms and revealed molecular signatures of different cancer types. Inclusion of a broader biomarker panel will further improve the assay sensitivity. This XNA-based technology can also be combined with NGS technology to cover a wide range of variants in many genes and so is applicable to the development of comprehensive DNA based assays for a wide range of cancer diagnostics.

## ACKNOWLEDGMENTS

Dr. Walter Bodmer is a senior scientific adviser of Medical Advisory Board, DiaCarta Inc. We would like to give our thanks to Elena Peletskaya, Farah Patell-Socha, Mauro Scimia, Jyoti Phatake, Ping Ding, Paul Z. Zhang, Simin Tang and Zhen Cui who involved this project at different time period and Paul Zhang for statistical analysis.

