## Supplemental for "A Novel Xenonucleic Acid Mediated Molecular Clamping Technology for Early Colorectal Cancer Screening"

**Supplementary Table 1**. Primer, Probe and XNA Sequences for ColoScape ™ Assay

| Gene Target | Primer/probe/XNA | 5'→3' sequence |
|---|---|---|
| APC E1309 | APC 1309TAQ-F B | GACGACACAGGAAGCAGATTCTGC |
| APC E1309 | APC 1309TAQ-R B | GCTCACAGGATCTTCAGCTGAT |
| APC E1309 | APC 1309Pr | TTCCAATCTTTTATTTCTGCTATGTG |
| APC E1309 | APC CS01 (APC1309 XNA) | CTGACCTAGTTCCAATCTTTTCTG |
| APC Q1367 | EAPC 1367F001 | TTCAGGAGCGAAATCTCCCTC |
| APC Q1367 | EAPC 1367R001 | TGAACATAGTGTTCAGGTGG |
| APC Q1367 | APC1367BHQnova | CAAAAGTGGTGCTTAGACACCCAAAAC |
| APC Q1367 | APC CS02 (APC 1367 XNA) | AGTGGTGCTCAGACACC |
| APC R1450 | APC3_1F002 | CCAGATAGCCCTGGACAAACCAT |
| APC R1450 | APC3_1R002 | CTTTTCAGCAGTAGGTGCTTTATTTTT |
| APC R1450 | APC1450_01 | AGGTACTTCTCaCTTGGTTTGA |
| APC R1450 | CS03.1 (APC 1450 XNA) | TAGGTACTTCTCGCTTGGTTTGA |
| APC R876 | APC876FT1 | TGAATGGCTGACACTTCTTCCATG |
| APC R876 | APC876RT1b | AGAAAATCCAGGAACTTCTTCAAGGAG |
| APC R876 | APC876Pr | TCTGGGCTGCAGTGGTGGAGATCTG |
| APC R876 | APC 876 XNA | GATCTGCAAACCTCGCTTTGA |
| CTNNB1 T41 | PB-CTNNB1-F | ACTCTGGAATCCATTCTGGTGCCA |
| CTNNB1 T41 | PB-CTNNB1-R | AGAAAATCCCTGTTCCCACTCATACA |
| CTNNB1 T41 | BCTM02SBHQ | AGGAAGAGGATGTGGATACCTCCCAAGTC |
| CTNNB1 T41 | CS05S (CTNNB1 41 XNA) | TGCCACTACCACAGCTCCT |
| CTNNB1 S45 | CS06S (CTNNB1 45 XNA) | AGCTCCTTCTCTGAGTG |
| KRAS G12 | KRASBioFP002_1 | AAGGCCTGCTGAAAATGACTGAA |
| KRAS G12 | KRASG12VBPR001_2 | GTTGGATCATATTCGTCCACAA |
| KRAS G12 | KRASCS02BHQnova | TCTGAATTAGCTGTATCGTCAAGGCACTCTTG |
| KRAS G12 | DPCK001C22 (KRAS 12 XNA) | CTACGCCACCAGCTCCAACTACCACA |
| KRAS G13 | C13F001_2 (KRAS G13 F primer) | ACTTGTGGTAGTTGGAGCTGGTG |
| KRAS G13 | DPCK002B1 (KRAS 13 XNA) | TCTTGCCTACGCCACCAGCTCCAAC |
| BRAF V600 | BRAF600EF003 | GGTGATTTTGGTCTAGCTACGGT |
| BRAF V600 | BRAFAZRP001_2 | CATCCACAAAATGGATCCAGACAACT |
| BRAF V600 | BRAF600P01BHQnova | CAAACTGATGGGACCCACTCCATCC |
| BRAF V600 | DPCBR001B1 (BRAFV600 XNA) | ATCGAGATTTCACTGTAGCTAGAG |
| ACTB | ACTBF3 | TCTGCCTTACAGATCATGTTTGAC |
| ACTB | ACTBR2 | CCAGAGGCGTACAGGGATAC |
| ACTB | ACTBPr2 | CCATGTACGTTGCTATCCAGGCTGA |

**Supplementary Table 2**. ColoScape $^{TM}$ Assay Panel

| ColoScape Assay Panel | FAM * | HEX | CFR610 (Texas Red or ROX) | CY5 (Internal control) |
|---|---|---|---|---|
| A | APC E1309 | APC Q1367 | | ACTB |
| B | APC R1450/R876 | KRAS G12 | CTNNB1 T41 | ACTB |
| C | BRAFV600E | KRAS G13 | CTNNB1 S45 | ACTB |

*Fam, HEX, CFR610 and CY5 used for probe labeling

**Supplementary Table 3**. Comparison of Single-plex and Multiplex for ColoScape $^{TM}$ Assay

| **Gene Target** | **1% mutant Ct singleplex** | **1% mutant Ct multiplex** | **ΔCt = Ct s - Ct m*** |
|---|---|---|---|
| Internal control | 28.60 | 28.40 | 0.20 |
| APC E1309 | 33.21 | 34.20 | -1.00 |
| APC Q1367 | 35.89 | 34.88 | 1.01 |
| APC R1450/R876 | 31.14 | 30.75 | 0.39 |
| CTNNB1 T41 | 32.61 | 32.54 | 0.07 |
| CTNNB1 S45 | 32.74 | 32.86 | -0.12 |
| KRAS G12 | 31.99 | 31.48 | 0.51 |
| KRAS G13 | 29.59 | 29.87 | -0.28 |
| BRAFV600E | 32.63 | 32.84 | -0.21 |

*Ct s: Ct from single-plex; Ct m: Ct from multiplex.

**Supplementary Table 4**. Summary of Assay Limit of Detection for gDNA Reference Standards

| Reference DNA | 5 ng DNA Input, ng/well | | | |
|---|---|---|---|---|
| Gene Target | Instrument | ABIQS5 | ABI 7500 Fast Dx | LC 480 II |
| | VAF% | % Correct Call | % Correct Call | % Correct Call |
| APC E1309 | 1% mutation | 100% | 100% | 100% |
| | 0.5% mutation | 100% | 100% | 100% |
| | 0.10% mutation | 25% | 90% | 55% |
| APC Q1367 | 1% mutation | 100% | 100% | 100% |
| | 0.5% mutation | 100% | 100% | 100% |
| | 0.10% mutation | 10% | 100% | 95% |
| APC R1450 | 1% mutation | 100% | 100% | 100% |
| | 0.5% mutation | 100% | 95% | 100% |
| | 0.10% mutation | 100% | 85% | 75% |
| CTNNB1 T41 | 1% mutation | 100% | 100% | 100% |
| | 0.5% mutation | 100% | 95% | 100% |
| | 0.10% mutation | 65% | 35% | 65% |
| CTNNB1 S45 | 1% mutation | 100% | 100% | 100% |
| | 0.5% mutation | 100% | 95% | 100% |
| | 0.10% mutation | 80% | 50% | 80% |
| KRAS G12 | 1% mutation | 100% | 100% | 100% |
| | 0.5% mutation | 100% | 100% | 100% |
| | 0.10% mutation | 95% | 95% | 95% |
| KRAS G13 | 1% mutation | 100% | 100% | 100% |
| | 0.5% mutation | 85% | 100% | 95% |
| | 0.10% mutation | 25% | 75% | 5% |
| BRAF V600 | 1% mutation | 100% | 100% | 100% |
| | 0.5% mutation | 100% | 100% | 100% |
| | 0.10% mutation | 100% | 100% | 100% |

**Supplementary Table 5**. Comparison of ColoScape $^{TM}$ and Sanger Sequencing for CRC FFPE

| Sample ID | Pathology Diagnosis | ColoScape $^{TM}$ | Sanger Sequencing* |
|---|---|---|---|
| WJZ4-C | (rectal) moderately differentiated adenocarcinoma | KRAS G13 | KRAS G13 |
| WJZ8-C | (sigmoid colon) differentiated adenocarcinoma | CTNNB1 T41 | CTNNB1 T41 |
| WJZ11-C | Right colon adenocarcinoma | KRAS G12 | KRAS G12 |
| WJZ9-C | Rectal paralysis | KRAS G12 | KRAS G12 |
| WJZ3-C | Colonic ulcer adenocarcinoma | KRAS G12 | KRAS G12 |
| WJZ17-C | Rectal adenocarcinoma, grade II | KRAS G12 | KRAS G12 |
| WJZ18-C | Sigmoid colonic adenocarcinoma, grade II | CTNNB1 T41 | CTNNB1 T41 |
| WJZ21-C | Right colon adenoma | Negative | Negative |
| WJZ23-C | Rectal tubular adenocarcinoma with multiple tubular adenomas of the colon | KRAS G12 | KRAS G12 |
| WJZ24-C | Rectal adenoma | Negative | Negative |
| WJZ27-C | Sigmoid colon adenoma | Negative | Negative |
| WJZ28-C | Rectal ulcer type moderately differentiated adenocarcinoma | KRAS G12 | KRAS G12 |
| WJZ29-C | Colon cancer ulcer type moderately differentiated adenocarcinoma | KRAS G13 | KRAS G13 |
| WJZ32-C | Rectal adenocarcinoma grade II-III with necrosis | KRAS G12 | KRAS G12 |
| WJZ33-C | Right colonic bulging adenocarcinoma grade II | APC R1450/R876 | APC R1450 WT/APC R876 |
| WJZ34-C | Colonic ulcer mucinous adenocarcinoma | KRAS G12 | KRAS G12 |
| WJZ35-C | Differentiated adenocarcinoma | KRAS G12 | KRAS G12 |
| WJZ36-C | Omental metastatic poorly differentiated carcinoma | KRAS G13 | KRAS G13 |
| WJZ38-C | (rectal) adenocarcinoma grade II | KRAS G13 | KRAS G13 |
| WJZ39-C | Medium and low differentiated adenocarcinoma in right colonic ulcer | KRAS G13 | KRAS G13 |
| WJZ40-C | Rectal prominence signet ring cell carcinoma | CTNNB1 S45 | CTNNB1 S45 |
| WJZ41-C | (rectal) medium-poorly differentiated adenocarcinoma | APC R1450/R876 | APC1450 WT/APCR R876 |
| WJZ42-C | colon, rectum) medium-poorly differentiated adenocarcinoma | KRAS G13 | KRAS G13 |
| WJZ43-C | (rectal) moderately differentiated adenocarcinoma | KRAS G13 | KRAS G13 |
| WJZ44-C | Rectal bulging tubular adenocarcinoma grade II | KRAS G12 | KRAS G12 |
| WJZ45-C | (rectal) adenocarcinoma grade II | CTNNB1 T41/KRAS G13/APC E1309/APC1367 | CTNNB1 T41/KRAS G13 |
| WJZ46-C | Rectal ulcer adenocarcinoma | KRAS G13 | KRAS G13 |

| | | | |
|---|---|---|---|
| WJZ52-C | (rectal sigmoid colon junction) adenocarcinoma | KRAS G12/APC E1309 | KRAS G12 |
| WJZ55-C | (rectal) moderately differentiated adenocarcinoma | KRAS G12 | KRAS G12 |
| WJZ56-C | Adenocarcinoma | KRAS G12 | KRAS G12 |
| WJZ57-C | (colon) adenocarcinoma | KRAS G13 | KRAS G13 |
| WJZ60-C | (rectal) adenocarcinoma grade II | KRAS G12 | KRAS G12 |
| WJZ62-C | Adenocarcinoma | KRASG12/APC E1309/APC Q1367 | KRAS G12 |
| WJZ63-C | colonic ulcer type moderately differentiated adenocarcinoma | APC E1309 | NA |
| WJZ67-C | Adenocarcinoma | APC R1450 /APC E1309/BRAFV600E | APC R1450/NA/BRAFV600E |
| WJZ68-C | Rectal adenocarcinoma | BRAFV600E | BRAFV600E |
| WJZ70-C | Rectal adenocarcinoma | KRAS G12 | KRAS G12 |
| WJZ71-C | Rectal adenocarcinoma | KRAS G12 | KRAS G12 |
| WJZ72-C | Rectal adenocarcinoma | KRAS G12 | KRAS G12 |
| WJZ73-C | Rectal adenoma | Negative | Negative |
| WJZ75-C | Colon adenocarcinoma | KRAS G12/APC E1309 | KRAS G12 |
| WJZ77-C | Rectal adenoma | Negative | NA |
| WJZ78-C | Adenocarcinoma | BRAFV600E | Poor sequencing data |
| WJZ79-C | Adenocarcinoma | KRAS G12 | KRAS G12 |
| WJZ80-C | (rectal) ulcerated adenocarcinoma grade II | Negative | NA |
| WJZ81-C | Rectal adenocarcinoma | KRAS G12 | KRAS G12 |
| WJZ84-C | Rectal adenocarcinoma | KRAS G12 | KRAS G12 |
| WJZ88-C | Adenocarcinoma | KRAS G12 | KRAS G12 |
| WJZ89-C | Rectal adenocarcinoma | KRAS G12 | KRAS G12 |
| WJZ90-C | Adenocarcinoma | KRAS G12/APC E1309 | KRAS G12 |
| WJZ91-C | Adenocarcinoma | KRAS G12 | KRAS G12 |
| WJZ92-C | Adenocarcinoma | KRAS G12 | KRAS G12 |
| WJZ93-C | Adenocarcinoma | KRAS G12 | KRAS G12 |
| WJZ95-C | Adenocarcinoma | KRAS G12 | KRAS G12 |
| WJZ96-C | Adenocarcinoma | KRAS G12 | KRAS G12 |
| WJZ97-C | Adenocarcinoma | KRAS G12 | KRAS G12 |
| WJZ98-C | Sigmoid colon cancer | KRAS G12 | KRAS G12 |
| WJZ99-C | (rectal) moderately differentiated adenoma | Negative | Negative |
| WJZ101-C | Adenocarcinoma | KRAS G12 | KRAS G12 |
| WJZ102-C | Colon cancer | BRAFV600E | BRAFV600E |
| WJZ103-C | Colon adenocarcinoma | KRAS G13 | KRAS G13 |

| | | | |
|---|---|---|---|
| WJZ104-C | Medium differentiated adenocarcinoma | BRAFV600E | BRAFV600E |
| WJZ106-C | Right colon adenoma | Negative | Negative |
| WJZ107-C | Colon adenocarcinoma | KRAS G12 | KRAS G12 |
| WJZ108-C | Rectal cancer | KRAS G12 | KRAS G12 |
| WJZ110-C | Rectal cancer | KRAS G13 | KRAS G13 |
| WJZ113-C | Adenocarcinoma | BRAFV600E | Poor sequencing data |
| WJZ114-C | Colon adenocarcinoma | KRAS G13 | KRAS G13 |
| WJZ116-C | Sigmoid colon adenocarcinoma | KRAS G13 | KRAS G13 |
| WJZ117-C | Rectal mucinous adenocarcinoma | KRAS G12 | KRAS G12 |
| WJZ118-C | Rectal adenocarcinoma | KRAS G12 | KRAS G12 |
| WJZ120-C | Sigmoid colon adenocarcinoma | BRAF V600E | Poor sequencing data |
| WJZ122-C | Rectal adenocarcinoma | BRAF V600E | Poor sequencing data |
| WJZ124-C | Sigmoid colon adenocarcinoma | KRAS G13 | KRAS G13 |
| WJZ125-C | Sigmoid colon adenocarcinoma | BRAF V600E | BRAF V600E |
| WJZ127-C | Rectal cancer | KRAS G12 | KRAS G12 |
| WJZ131-C | Rectal adenocarcinoma | KRAS G12 | KRAS G12 |
| WJZ132-C | Adenocarcinoma | KRAS G12 | KRAS G12 |
| WJZ133-C | Adenocarcinoma | KRAS G12 | KRAS G12 |
| WJZ134-C | Right colon adenocarcinoma | BRAF V600E | Poor sequencing data |
| WJZ135-C | Colon adenocarcinoma | KRAS G12 | KRAS G12 |
| WJZ136-C | Colon adenocarcinoma | KRAS G12 | KRAS G12 |
| WJZ138-C | Sigmoid colon cancer | KRAS G12 | KRAS G12 |
| WJZ140-C | Rectal mucinous adenocarcinoma | KRAS G13 | KRAS G13 |
| WJZ142-C | sigmoid colon) ulcerated tubular adenocarcinoma | KRAS G12 | KRAS G12 |
| WJZ143-C | Colonic bulging adenocarcinoma grade II | KRAS G13 | KRAS G13 |
| WJZ144-C | Sigmoid colon adenocarcinoma | KRAS G13 | KRAS G13 |
| WJZ145-C | Colon adenocarcinoma | KRAS G12 | KRAS G12 |
| WJZ146-C | Rectal adenocarcinoma | BRAF V600E | BRAF V600E |
| WJZ147-C | Colon adenocarcinoma | KRAS G12 | KRAS G12 |
| WJZ148-C | Colon adenocarcinoma | KRAS G12 | KRAS G12 |
| WJZ149-C | Colon adenocarcinoma | KRAS G12 | KRAS G12 |
| WJZ150-C | Sigmoid colon cancer | BRAF V600E | Poor sequencing data |
| WJZ151-C | Colon cancer | KRAS G12 | KRAS G12 |
| WJZ152-C | Colon cancer | KRAS G13 | KRAS G13 |
| WJZ153-C | Colon adenoma | Negative | Negative |
| WJZ154-C | Sigmoid colon adenocarcinoma | KRAS G12 | KRAS G12 |

*qPCR amplicons were sequenced by Sanger Sequence method.

**Supplementary Table 6**. Sensitivity of KRAS G12 Variant Mutation Detection

| KRAS G12 mutation | G12A | G12R | G12D | G12C | G12S | G12V |
|---|---|---|---|---|---|---|
| LOD (VAF%) | 0.2% | 2% | 0.2% | 2.5% | 2% | 0.2% |

*gDNA reference was used
